# Knocking out CD70 rescues CD70-specific nanoCAR T cells from antigen induced exhaustion

**DOI:** 10.1101/2023.01.22.523482

**Authors:** Stijn De Munter, Juliane Buhl, Laurenz De Cock, Alexander Van Parys, Willem Daneels, Eva Pascal, Lucas Deseins, Joline Ingels, Glenn Goetgeluk, Lore Billiet, Melissa Pille, Niels Vandamme, Jo Van Dorpe, Fritz Offner, Erik Depla, Jan Tavernier, Tessa Kerre, Jarno Drost, Bart Vandekerckhove

## Abstract

CD70 is an attractive target for chimeric antigen receptor (CAR) T cell therapy as treatment for both solid and liquid malignancies. However, functionality of CD70-specific CARs is only modest. Here, we optimized a CD70-specific VHH based CAR (nanoCAR). We evaluated the nanoCARs in clinically relevant models *in vitro*, using co-cultures of CD70-specific nanoCAR T cells with malignant rhabdoid tumor organoids, and *in vivo* by using a diffuse large B cell lymphoma (DLBCL) patient-derived xenograft (PDX) model. Whereas the nanoCAR T cells were highly efficient in organoid co-cultures, they showed only modest efficacy in the PDX model. Knocking out CD70 expression by the nanoCAR T cells resulted in dramatically enhanced functionality in the PDX model, suggesting that endogenous CD70 interaction with the nanoCAR induces exhaustion. Through single-cell transcriptomics, we obtained evidence that CD70KO CD70-specific nanoCAR T cells are protected from antigen induced exhaustion. Our data shows that CARs targeted to endogenous T cell antigens, negatively affect CAR T cell functionality by inducing an exhausted state which can be overcome by knocking out the specific target, in this case CD70.

## Introduction

Adoptive transfer of chimeric antigen receptor (CAR) T cells has demonstrated great potential for the treatment of various hematological malignancies^1–7^. While response rates are high, relapses still occur due to the emergence of antigen escape variants or due to the lack of long term *in vivo* persistence of the CAR T cells^1,4,5,8–10^. Furthermore, the efficacy in solid tumors has yet to be demonstrated as the tumor suppressive microenvironment, homing inside the tumor, tumor heterogeneity and the lack of optimal target antigens may compromise their efficacy^11–13^.

CD70 is an attractive and broadly applicable target antigen. It is a tumor necrosis factor (TNF) superfamily member that acts as co-stimulatory ligand for CD27. CD70 has a broad expression pattern on multiple solid and liquid malignancies such as clear cell renal cell carcinoma, pancreatic cancer and some hematological malignancies such acute myeloid leukemia (AML) and diffuse large B cell lymphoma. Furthermore, expression has been reported on both primary and metastatic tumor samples. Additionally, CD70 expression is associated with a poor prognosis and a decreased survival^14–16^. CD70 has been extensively studied in the context of AML, for which it has been shown by Riether *et al*. that the ligand-receptor pair CD70/CD27 is expressed on AML blasts and AML stem/progenitor cells. This results in cell-autonomous CD70/CD27 signaling which propagates the disease through activation of a stem cell gene expression program, Wnt pathway activation and promotion of symmetric cell divisions and proliferation^17^. Recently, cusatuzumab, an antibody targeting CD70, showed impressive response rates in a phase I trial for AML patients unfit for traditional intensive-induction chemotherapy^18,19^. CD70 is not expressed in healthy tissues nor on hematopoietic stem cells but is transiently upregulated on activated immune cells such as B cells, T cells and dendritic cells ^15^. In activated T cells, it provides survival signals and influences memory formation. CD70 specific CAR T cells were reported to inhibit anti-viral immune responses^20^.

CARs against CD70 have been developed by several groups using different strategies. A first strategy utilized a ligand-based targeting approach, the CD27 protein fused to the CD3ζ domain (flCD27-ζ), and showed efficacy towards multiple different B cell malignancies^21^. This design was further optimized by fusing extracellular CD27 with intracellular 4_1BB (trCD27-4_1BB:ζ) resulting in both a higher *in vivo* and *in vitro* functionality^22^. This construct is subject of an ongoing phase I/II clinical trial (NCT02830724). A last optimization was performed by Leick and colleagues: in a two way approach they increased the functional avidity and thus functionality of the trCD27-4_1BB:ζ CAR^23^. Next to the ligand-based targeting approach, a second strategy uses single chain variable fragments (scFvs) as target antigen recognition domains. A study by Sauer *et al*. showed that the original flCD27-ζ CAR outperformed a panel of scFv based CARs^20^. In a more recent study, multiple scFvs all targeting CD70 were evaluated. Two distinct classes of scFvs were defined based on their CD70-specific binding epitope. When used in CARs, these resulted in CAR T cells with different memory phenotypes, activation statuses and cytotoxic activity^24^.

While CD70 is an attractive target antigen due to its expression on multiple different tumor types, CD70 expression on malignant cells both solid as liquid, is lower and less uniform compared to the expression of CD19 in B cell malignancies. Since CAR T cell activity is dependent on antigen load, it could be necessary to increase the expression level of CD70 on the cell membrane^1,8,25–27^. Indeed, it has been proven in a pre-clinical study that CD70-specific CAR T cells have a higher efficacy when combined with the hypomethylating agent azacitidine which acts as a nucleoside analog, inhibiting DNA methyltransferase and decreasing methylation of the CD70 promotor, resulting in an increased surface expression of CD70^23^. The use of azacitidine has also been assessed in a clinical trial using the CD70 specific monoclonal antibody cusatuzumab^19^. A similar approach has also shown its merit in a preclinical model of a CD22-targetting CAR. Here, bryostatin 1, a macrocyclic lactone that acts as modulator of kinase C, was used to increase the antigen density of CD22 on leukemic cell lines, resulting in higher CD22 expression and a higher efficacy of CD22-specific CAR T cells^26^.

While scFvs are commonly used as targeting domain in CAR T cells, also other antigen binding moieties can be used. We have previously shown that nanobodies™, the variable regions of heavy chain only antibodies (VHH), can be used as antigen recognition domains in CARs. We termed these CARs as nanoCARs^28,29^ Due to their monomeric structure they do not require a conversion from immunoglobulin (Ig) towards single chain variable fragment (scFv). Furthermore, they have a high solubility and lack aggregative properties which reduce the likelyhood of CAR clustering and tonic signaling^30^. Lastly, due to their smaller size and high sequence homology with the human VH3 gene family, they have a lower immunogenicity as compared to murine scFvs^31,32^. If needed, they can be easily humanized^33^.

In this study, we optimized a CD70-specific nanoCAR. First, we generated multiple nanoCAR constructs which differ in co-stimulatory moiety. The lead nanoCAR was selected based on the best *in vivo* functionality and persistence. Next, we evaluated the lead nanoCAR *in vitro* and *in vivo* against patient derived tumor cells in clinically relevant model systems. We observed complete elimination of organoid cultures of malignant rhabdoid tumors at very low effector to target (E:T) ratios. However, when tested in a DLBCL patient-derived xenograft (PDX) model, only a modest efficacy could be observed. Using single-cell transcriptomics, we could show that tumor-infiltrated CD70-specific nanoCAR T cells acquire an exhausted phenotype during activation resulting in less potent nanoCAR T cells. *CD70* gene deletion in the nanoCAR T cells dramatically altered the T cell transcriptome and improved *in vivo* functionality. Our data shows that CARs targeted to T cell antigens could negatively alter CAR T cell efficacy by inducing an exhausted state and that this could be overcome by knocking out the specific target, in this case CD70.

## Results

### JAK-STAT activation does not enhance CD70-specific nanoCAR functionality

Out of a library of 33 llama VHH, the VHH with the highest affinity for CD70 was selected. It was reported that addition of STAT3 and STAT5 docking sites to the cytoplasmic tail of the CAR improves activity against solid tumors^34^. Therefore, we generated a panel of four CD70-specific nanoCAR constructs with different transmembrane and/or intracellular domains. Two second generation nanoCARs were generated: i) nanoCAR with CD8α as spacer and transmembrane domain and 4_1BB as co-stimulatory domain (70-4-1BB:ζ) and ii) nanoCAR with CD28 as spacer, transmembrane domain and co-stimulatory domain (70-28:ζ) (Figure 1A). In addition, we incorporated a truncated cytoplasmic domain of the interleukin −2 (IL-2) receptor β between the cytoplasmic domains of 4_1BB (70-4-1BB:β:ζ*) or CD28(70-28:β:ζ*) and CD3ζ for STAT5 recruitment, and added a YXXQ motif at the C-terminus of CD3ζ for STAT3 recruitment, as described previously^34^ (Figure 1A). NanoCAR T cells were generated by retroviral transduction. After staining with a VHH specific antibody, we could validate expression of all the nanoCAR constructs on the cell surface (Figure 1B). However, nanoCARs containing the truncated IL-2 receptor β chain showed a lower mean fluorescent intensity (MFI) (Figure 1C). Furthermore, 70-4_1BB:β:ζ* showed a consistently lower transduction efficiency as compared to the other three nanoCAR constructs (Figure 1D). We tested the functionality of the different nanoCAR constructs *in vitro* against the CD70^−^ cell line Nalm6 and the CD70^+^ cell line SKOV3 (Figure S1A). IFN-γ and IL-2 were produced upon incubation with the CD70^+^ cell line, while there was no production when the nanoCAR T cells were incubated with Nalm6 (Figure 1E). All nanoCAR T cells showed cytokine production and the highest percentage of IL-2^+^ cells could be observed in nanoCARs containing the CD28 co-stimulatory domain. Interestingly, 70-4_1BB:β:ζ* nanoCAR T cells showed an overall lower cytokine production profile for both the CD4^+^ and the CD8^+^ T cells, suggesting a potential lower functionality. To evaluate the *in vivo* antitumoral effect of the CD70-specific nanoCAR T cells, a SKOV3 murine xenograft model was used, in which NSG mice were subcutaneously injected with the CD70^+^ ovarian cancer cell line SKOV3 (Figure 1F). Different concentrations of CD70-specific nanoCAR T cells were injected and tumor growth was measured by caliper. At the highest nanoCAR T cell doses, all four nanoCARs showed similar efficacy (Figure S1B-E). When the dose was lowered, the 70-4_1BB:β:ζ* nanoCAR, combining both 4_1BB as JAK-STAT signaling, was no longer capable of complete tumor eradication nor of tumor control (Figure S1D-E). In addition, at the lowest dose, only second generation nanoCARs showed similar high efficacy (Figure 1G-I). Antitumoral responses were long lasting for both second generation nanoCARs but not for the 70-28:β:ζ* incorporating JAK-STAT signaling (Figure 1G-I). At day 63 post nanoCAR T cell injection, we assessed the presence of nanoCAR T cells in the spleen of all treated mice (Figure 1H). We observed the highest frequency of nanoCAR^+^ T cells in mice that were treated with a standard 4_1BB second generation nanoCAR, and the lowest frequency in mice treated with the CD28 nanoCAR containing the domains for JAK-STAT signaling. These data show that JAK-STAT signaling did not enhance nanoCAR T cell functionality in this model and resulted in a less efficient nanoCAR T cell product. The 70-4_1BB:ζ second generation nanoCAR outperformed all other nanoCAR T cell constructs in terms of *in vivo* efficacy and persistence. As a result, we selected the 70-4-1BB:ζ CAR for further optimization.

**Figure 1:**
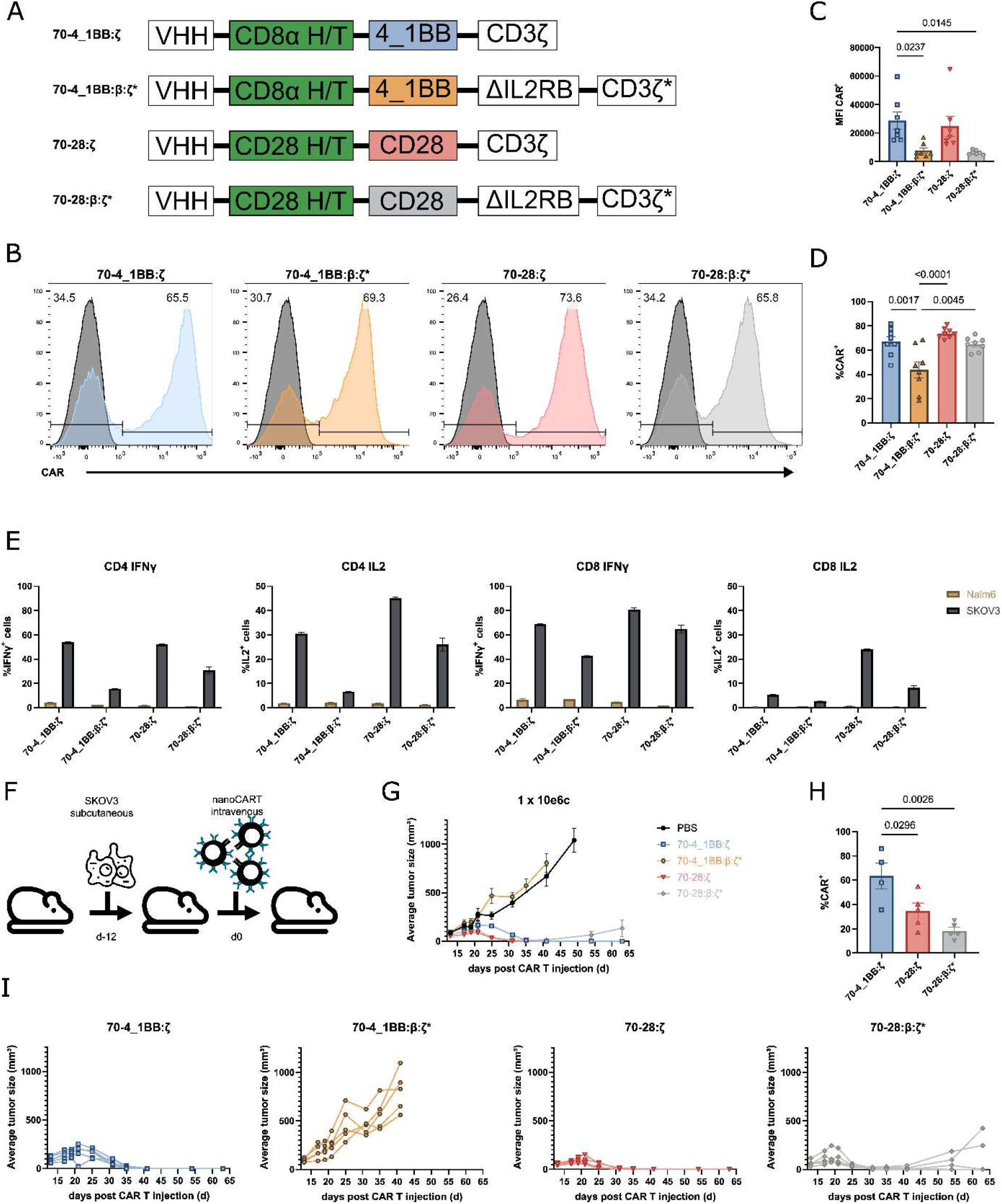
STAT signaling does not affect CD70-specific CAR T cell functionality. **(A)** Design of the different CAR T cell constructs used. **(B)** Representative histograms of CD70 CAR expression levels after transduction of activated T cells, nine days post transduction. **(C)** MFI of CAR positive population. **(D)** The frequency of CAR positive T cells of eight different donors. **(E)** Cytokine production by CD70-specific CAR T cells after stimulation with Nalm6 (CD70^−^) and SKOV3 (CD70^+^), determined by intracellular staining for IFNγ and IL2. **(F)** The experimental setup of the SKOV3 xenograft model. CD70-specific T cells (1×10e6) were injected intravenously twelve days after subcutaneous engraftment of SKOV3 cells (1×10e6) (n=5 for each treatment group). **(G)** SKOV3 tumor growth after injection of CAR T cells. Tumor growth is shown as average tumor size, error bars indicate SEM. **(H)** Tumor growth for each individual mouse. **(I)** Percentage of CAR^+^ cells in the CD45^+^CD3^+^ in the spleen at day 63 post CAR T cell injection as assessed by flow cytometry. Mean values are shown, error bars indicate SEM.

### Malignant rhabdoid tumor-derived organoids are efficiently killed by CD70-specific nanoCAR T cells

While cell lines are ideal for fundamental research, they are less relevant for translational research due to loss of heterogeneity (e.g. target antigen expression) and divergence from the original tumor present in the patient. To overcome this hurdle and show the potential of the CD70-specific nanoCAR, we assessed the efficacy of CD70 nanoCAR T cells in CD70 expressing malignant organoid structures. These structures are derived from patient malignant tissue and retain the tumor heterogeneity observed in patients. First, we analyzed CD70 mRNA levels in Wilms tumor and malignant rhabdoid tumor (MRT) patient samples using data from Calandrini *et al*.^35^. While we observed no CD70 expression in primary Wilms tumor, mRNA was expressed in all MRT samples to a variable degree (Figure 2A). Next, CD70 mRNA levels were quantified in organoid lines derived from normal kidney, Wilms tumor and MRT^35^. In line with the tissues, normal kidney and Wilms tumor organoid lines showed no expression of CD70 while CD70 was expressed to variable degrees in MRT organoid lines, similar to the original tumor (Figure 2B). CD70 membrane expression was further validated in four MRT organoid lines using flow cytometry. This revealed that three out of the four lines display low to high CD70 membrane expression (Figure 2C). One line (JD041T) did not show any CD70 expression and was taken as negative control in all subsequent experiments. NanoCAR T cells were incubated with different organoid lines at multiple target to effector ratios. A CD20-specific nanoCAR was used as negative control^29^ (Figure 2D). Viable MRT cells were measured based on luciferase activity over time. The CD70-specific nanoCAR T cells induced strong and specific tumor lysis of all three antigen positive organoid lines even at target to effector ratios of 100:1 (Figure 2E). The weakest cytotoxicity was seen after culture with the organoid line 60T2 when compared to the other organoid lines, although CD70 expression was high in these organoids (Figure 2D-E). In addition, staining of the activation marker CD137 on nanoCAR T cells showed strong expression after 20 hours of incubation with the organoid lines (Figure 2F). Expression of CD137 was in line with the killing efficiency towards the different organoid lines, high expression correlated with high nanoCAR T cell efficacy. The CD70^−^ organoid line JD041T was unaffected by the CD70-specific nanoCAR T cells. Some background nonspecific killing activity was observed by both the CD70 as the CD20 CAR T cells, probably caused by allogeneic reactivity (Figure 2D). Lastly, we assessed the motility of the CAR T cells in the different organoid structures. We noticed interactions between the CD70-specific nanoCAR T cells with organoid cells (Movie 1-3). These interactions resulted in killing of the CD70^+^ organoid cells and proliferation of the CD70-specific nanoCAR T cells (Figure 2G).

**Figure 2:**
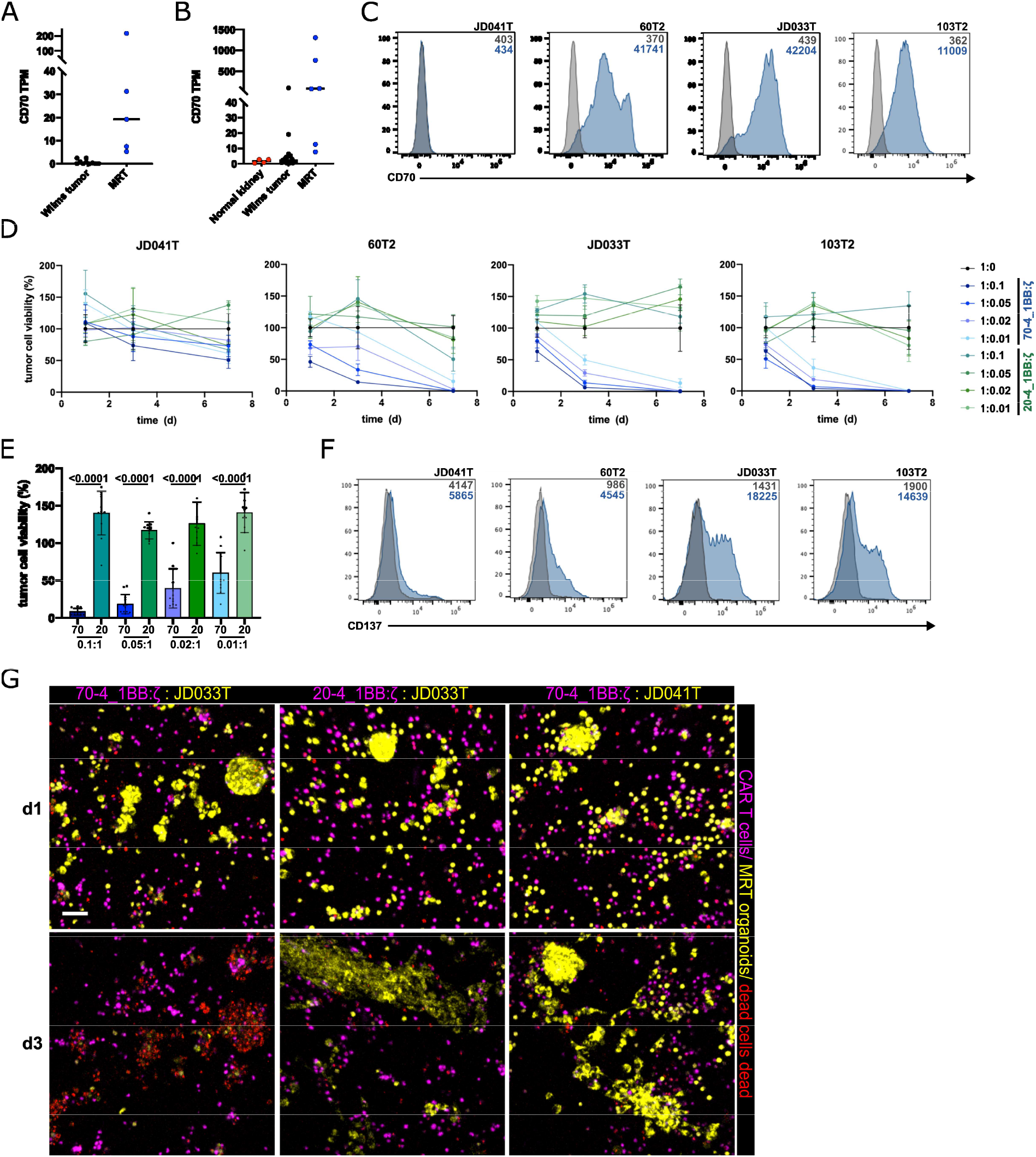
CD70 specific CAR T cells are reactive towards CD70 MRT organoids. **(A)** Normalized CD70 transcript per million (TPM) values in Wilms tumor and MRT patient samples. **(B)** Normalized CD70 transcript per million (TPM) values in normal kidney, Wilms tumor, and MRT organoid lines. **(C)** Flow cytometric analysis of CD70 expression in 4 MRT organoid lines (number indicate MFI, blue = CD70 expression, grey = unstained). Results are representative of at least n=3 experiments. **(D)** Remaining luciferase activity in 4 MRT organoid lines after co-culture with 70-4_1BB:ζ or 20-4_1BB:ζ for 1, 3 and 7 days in the depicted target to effector ratios. Values are normalized to MRT organoids in culture alone. Mean ± SD of n=4 technical replicates. **(E)** Remaining luciferase activity in 3 CD70 positive MRT organoid lines at day 3 after co-culture with 70-4_1BB:ζ or 20-4_1BB:ζ in the depicted effector to target ratios. Mean ± SD of n=3 biological replicates. Statistical significance was analyzed by unpaired t-test. **(F)** Flow cytometry analysis of CD137 expression on 70-4_1BB:ζ (blue) and 20-4_1BB:ζ (grey) after co-culture with the depicted MRT organoid lines for 20 hours in an effector to target cell ratio of 0.1:1 (number indicate MFI, blue = CD137 expression, grey = unstained). **(G)** Immunofluorescence three-dimensional imaging of MRT organoid lines (yellow) JD033T (CD70 pos.) in co-culture with 70-4_1BB:ζ and 20-4_1BB:ζ (pink) as well as JD041T (CD70 neg.) with 70-4_1BB:ζ in a 1:1 E:T ratio on day 1 and 3. Dead cells are immunolabeled in red. Maximum intensity projections are shown. Scale bars: 50 μm.

### CD70-specific nanoCAR T cells show efficacy in primary DLBCL PDX model

Next, we explored the *in vivo* functionality of CD70-specific nanoCARs using clinical relevant material. DLBCL were reported to express CD70^15^. We selected a DLBCL PDX that expressed low levels of CD70 (Figure 3A). Fresh PDX cells were injected on day −28 and treated with 1 × 10e6 nanoCAR T cells on day 0. Treatment was evaluated by daily body weight measurements (Figure 3B). Mice treated with CD70-specific nanoCAR T cells performed significantly better compared to PBS control, although the antitumor effect was modest (Figure 3C). Although mice were succumbing to disease, we still observed both nanoCAR T cells and DLBCL cells 40 days post nanoCAR T cell injection in the liver, an organ which is consistently infiltrated in this DLBCL PDX model (Figure 3D), suggesting loss of nanoCAR T cell efficacy. Since we selected a DLBCL PDX with low CD70 expression, we speculated whether antigen escape was causing treatment failure. Antigen escape mutations are a common cause of CAR T cell failure^10^. CD70 expression on blood DLBCL cells decreased over time in nanoCAR treated mice. This reduction was observed starting at day 7 post nanoCAR T cell treatment (Figure 3E). At day 21 post nanoCAR T cell injection, virtually no CD70 expression could be detected in the mice treated with nanoCAR T cells, suggesting antigen escape (Figure 3E). However, sequencing of the CD70 cDNA showed no mutations (not shown). Next, the CD70^−^ DLBCL cells of CAR T treated mice were isolated and injected in untreated NSG mice. After 28 days, mice were sacrificed and CD70 expression was determined by flow cytometry. We observed CD70 membrane re-expression comparable to the expression by DLBCL cells harvested from untreated mice (Figure 3F). This suggests that the DLBCL cells in CAR T treated mice expressed *CD70* but membrane expression was downmodulated by interaction with the CAR receptor as previously described^36^.

**Figure 3:**
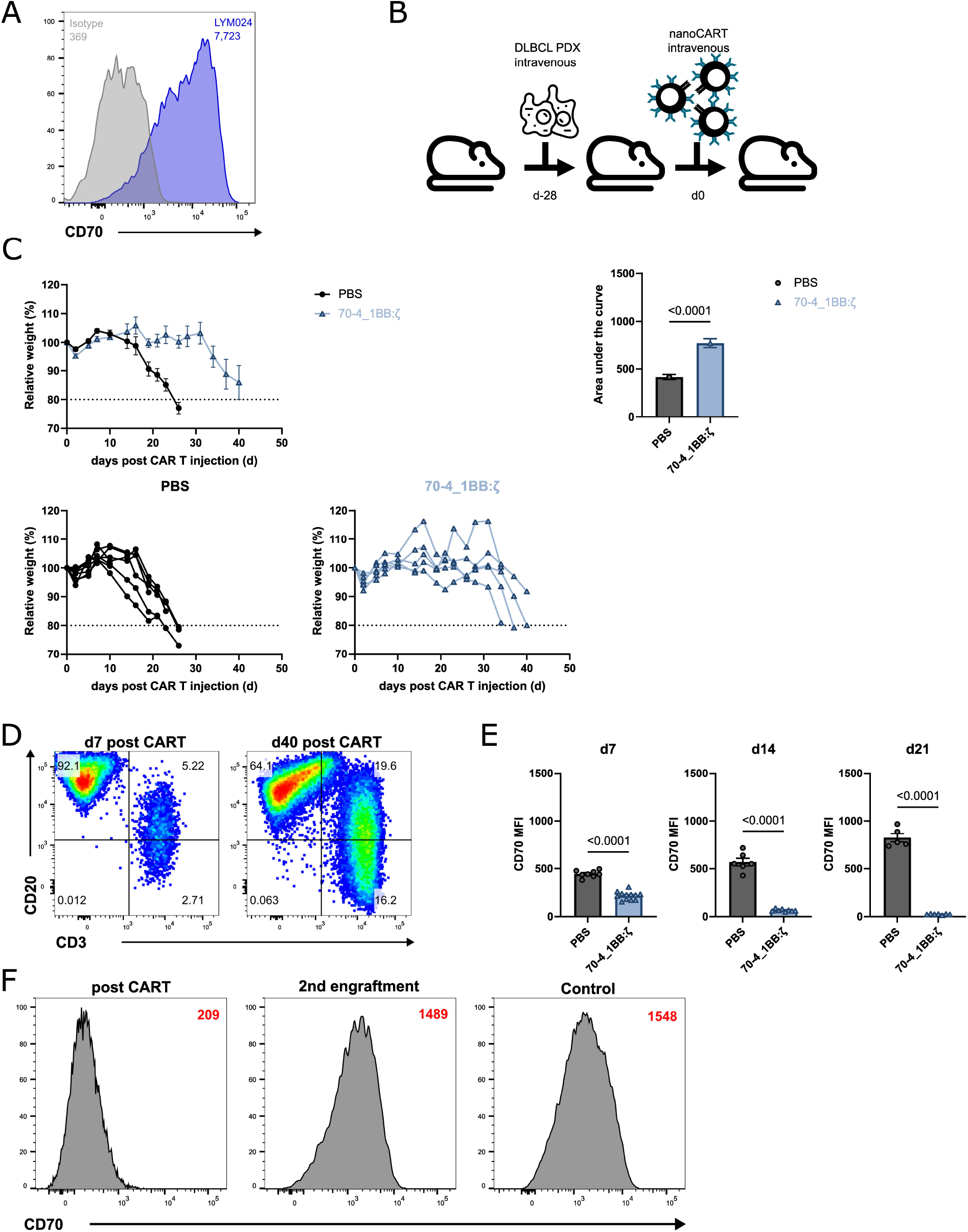
CD70 nanoCAR T cells are functional in a clinically relevant DLBCL PDX model. **(A)** C70 expression relative to isotype control of the LYM024 DLBCL PDX. **(B)** The experimental setup of the DLBCL PDX model. CD70 CAR T cells (1×10e6) were injected intravenously 28 days after intravenous injection of DLBCL cells (1×10e6) (n=5 for each treatment group). **(C)** Relative weight after CAR T cell injection. Weight at the start of the experiment was 100%. Area under the curve was calculated and significance was tested. **(D)** Presence of T cells (CD3^+^CD20^+/-^) and DLBCL cells (CD3^−^CD20^+^) at day seven and day 42 post nanoCAR T cell injection in the liver. **(E)** CD70 expression levels assessed by flow cytometry on circulating DLBCL cells at different time points post CAR T cell injection. Mean values are shown, error bars indicate SEM. **(F)** CD70 expression on DLBCL cells post CAR T cell treatment, after second engraftment and control. MFI is each time indicated.

### CD70 knock out in CD70-specific nanoCAR T cells reverses dysfunctionality

Since antigen escape mutations were not causing CD70-specific nanoCAR T cell treatment failure, we speculated whether CD70 expression by CAR T cells could influence their functionality. It is known that CD70 is expressed on activated T cells, however, CD70 expression on activated CAR T cells has not been reported^20,37^. CD70-specific nanoCAR T cells were consistently negative for CD70 membrane expression, possibly caused by CD70-CAR interaction (not shown). We therefore assayed nanoCAR T cells with a different specificity (CD20) and stimulated these with CD20^+^ Raji cells (Figure 4A). Unstimulated CD20-specific nanoCAR T cells already expressed CD70 at high levels. By 24 hours after stimulation, almost all of the cells expressed CD70. Expression further increased with the highest levels of CD70^+^ cells observed at 48 hours post-incubation. Furthermore, there was a robust raise of CD70 MFI up to 48 hours after stimulation (Figure 4A). Since CD70 was expressed and strongly upregulated upon activation of CD20-specific nanoCAR T cells, we argued that endogenous CD70 expression on CD70-specific nanoCAR T cells may be causing nanoCAR T cell dysfunctionality due to interaction with the nanoCAR either at the intracellular level and/or on the membrane of the T cells. We first validated CD70 knock out (CD70KO) generation in CD20 nanoCAR T cells. No CD70 expression could be observed in unstimulated and stimulated CD20 nanoCAR T cells in which CD70 was knocked out, showing the efficiency of *CD70* targeted gRNA and CRISPR/Cas9 technology (Figure 4A). Next, CD70KO CD70-specific nanoCAR T cells were generated. The functionality of CD70WT and CD70KO was assessed in a long-term *in vitro* assay. CD70 wild type (WT) and CD70KO CD70-specific nanoCAR T cells were stimulated with Raji cells every two days and cell numbers were determined. CD70KO nanoCAR T cells showed a greater potential to control Raji cell outgrowth than CD70WT nanoCAR T cells (Figure 4B). Furthermore, CD70KO CAR T cells showed a higher survival rate following antigen exposure compared to CD70WT CAR-T cells (Figure 4B). Of note, a similar enhancing effect of CD70KO was not observed for CD20 specific nanoCAR T cells (Figure S2A). To determine the *in vivo* effect of CD70KO in CD70-specific nanoCAR T cells, we injected NSG mice intravenously with CD70^+^ Raji cells followed by a low dose CD70WT or CD70KO nanoCAR T cells (Figure 4C). Mice injected with CD70WT nanoCAR T cells were unable to control Raji outgrowth and succumbed to disease similar to PBS injected animals. In contrast, CD70KO nanoCAR T cells induced improved tumor control and prolonged survival compared to the control groups indicating a robust increase in nanoCAR T cell efficacy (Figure 4D-G). In the animals treated with CD70KO nanoCAR T cells that did show tumor outgrowth, tumor cells were confined to the brain which suggests that the blood brain barrier may hamper nanoCAR T cell efficacy against brain metastases (Figure 4G).

**Figure 4:**
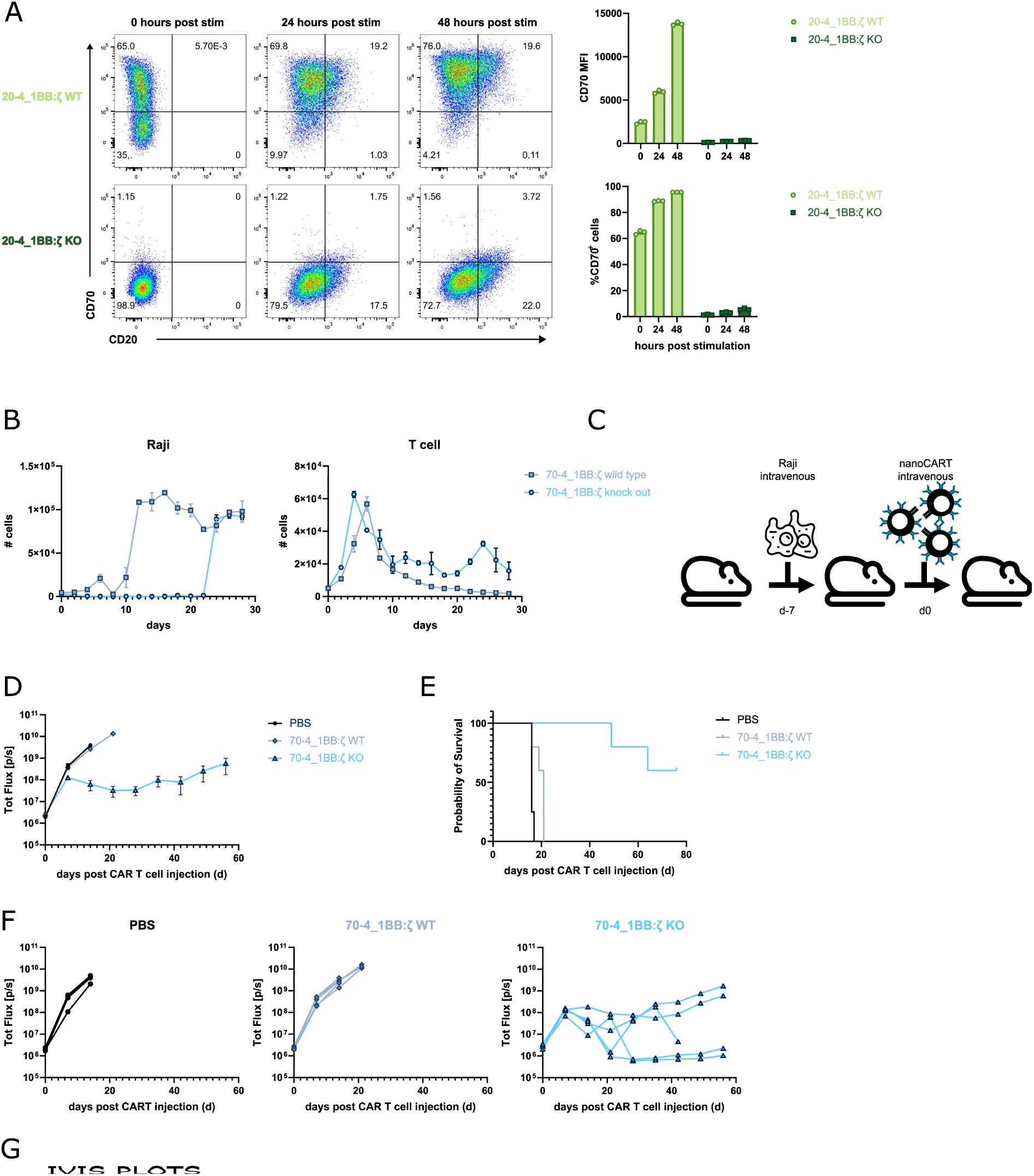
CD70 KO on CD70 specific nanoCAR T cells increases nanoCAR T cell functionality. **(A)** CD20 specific nanoCAR T cells, either WT or CD70 KO, were incubated with CD20^+^CD70^−^ Raji cells for zero, 24 or 48 hours. CD70 expression on the nanoCAR T cells was determined by flow cytometry. **(B)** CD70 specific nanoCAR T cells either WT or KO were incubated with WT Raji cells. Every two days the amount of cells were determined and the remaining wells were stimulated with fresh Raji cells. Median values are shown, error bars indicate SEM. **(C)** The experimental setup of the Raji xenograft model. CD70 nanoCAR T cells (0.75×10e6) were injected intravenously 7 days after intravenous injection of Raji cells (0.5×10e6). **(D)** Quantification of total flux (photons/s) in the experimental groups at the indicated time points. Points represent mean, error bars indicate SEM. **(E)** Kaplan-Meier survival curves for the different treatment groups. **(F)** Quantification of total flux (photons/s) for each mouse in the experimental groups at the indicated time points. **(G)** Bioluminescence of Raji xenografts over time in the treatment groups.

Next, we compared the CD70KO and CD70WT CD70-specific nanoCAR T cells in the patient relevant DLBCL PDX model following the same workflow as described earlier (Figure 3B). The weight of all mice dropped starting from day 5 post CAR T cell injection (Figure 5A-B). In mice treated with PBS, all mice had to be sacrificed around day 20 due to severe weight loss. Mice treated with CD70WT nanoCAR T cells experienced prolonged weight loss. Conversely, mice treated with CD70KO nanoCAR T cells showed an initial weight drop, probably caused by extensive nanoCAR T cell activation and the induction of cytokine release syndrome, but gained weight steadily afterwards and fully recovered. Weekly blood monitoring by flow cytometry revealed initial tumor control in the CD70WT nanoCAR T cell treated mice (Figure 5C-D). However, at day 21 post nanoCAR T cell injection, this control was lost. In contrast to the CD70WT nanoCAR T cell treated mice, we could not detect any tumor cells around day 21 in mice treated with CD70KO nanoCAR T cells. Tumor control coincided with expansion of nanoCAR T cells in the blood. This massive expansion of CD70KO nanoCAR T cells was not detected in mice treated with CD70WT nanoCAR T cells (Figure 5C,E).

**Figure 5:**
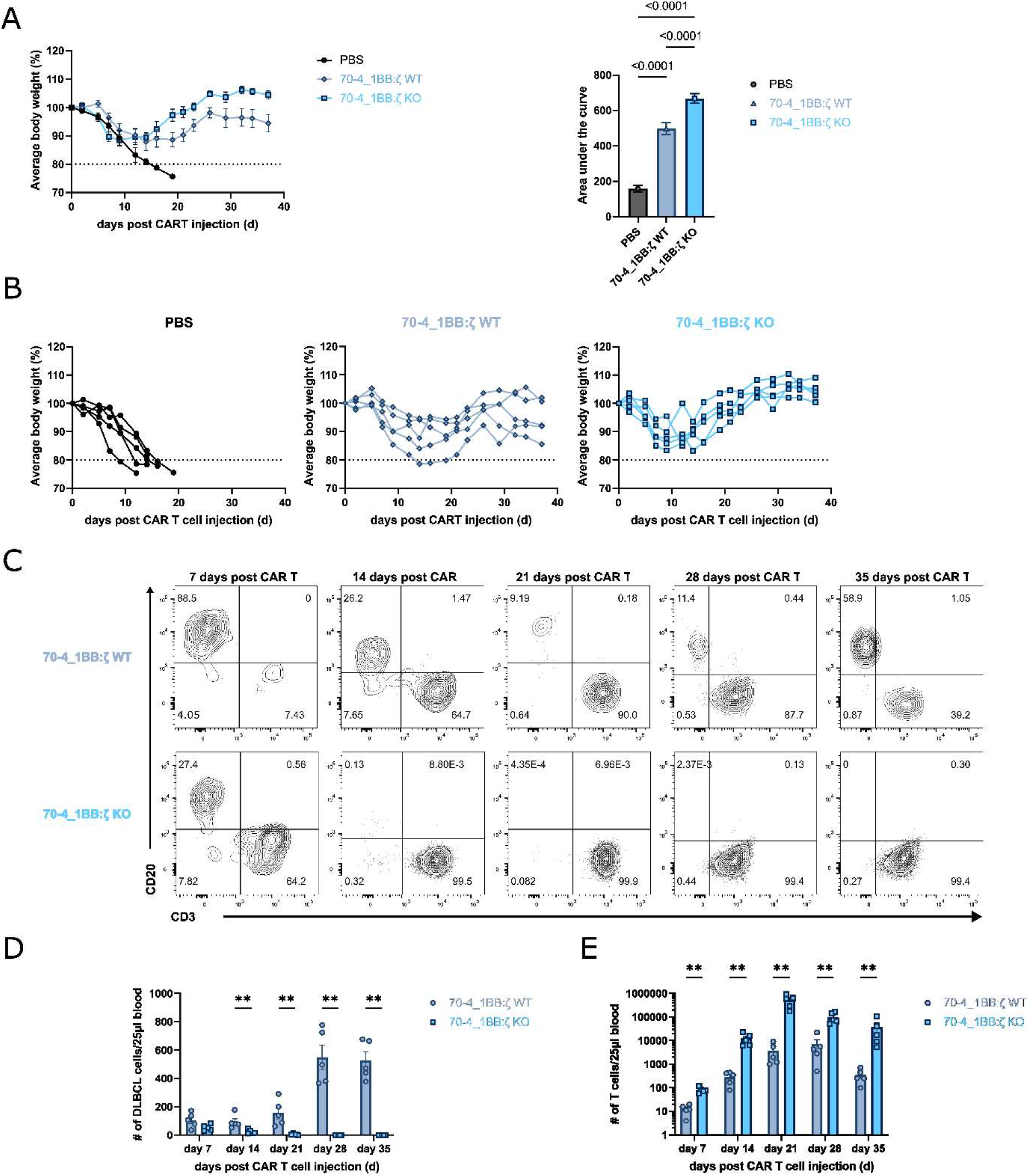
CD70 KO nanoCAR T cells eradicate DLBCL in a PDX model. **(A)** Relative weight after nanoCAR T cell injection. Weight at the start of the experiment was 100%. Mean number is shown, error bars indicate SEM. Area under the curve was calculated and significance was tested. **(B)** Relative body weight for individual mice for each treatment group. **(C)** Representative DLBCL and T cell expansion in blood at the indicated time points analyzed by flow cytometry. **(D)** Quantification of DLBCL and T cells in 25 μl blood for each treatment group. Mean number of cells is shown, error bars indicate SEM.

### CD70 KO nanoCAR T cells are protected from CAR induced exhaustion

To decipher the mechanism underlying the functional superiority of CD70KO CD70 specific nanoCAR T cells compared to their CD70WT counterparts, we analyzed the nanoCAR T cell tumor infiltrate seven days after injection using single-cell transcriptomics. Mice were injected with Raji cells and seven days later injected with either CD70WT nanoCAR T cells or C70KO nanoCAR T cells. Another seven days later, mice were sacrificed, and bone marrow was isolated (Figure 6A). At that time, Raji cells were present in both conditions (Figure S3A). The resulting cell suspension was labelled with hashtags and sorted for nanoCAR T cells. Sorted cells were pooled in roughly equal numbers and subsequently processed in a single run (Figure 6A). Uniform manifold approximation and projection (UMAP) analysis revealed twelve clusters among the sorted nanoCAR T cells (Figure 6B, S3B). Cluster 11 (Table 1) was excluded from further analysis since it was solely composed of mouse cells, caused by impurities from the sorting process. Substantial heterogeneity was observed in the contribution of CD70WT and CD70KO cells to each cluster. Cluster 0, 1, 4 and 9 were mainly composed of CD70KO nanoCAR T cells while these were largely excluded from cluster 2, 3, 8 and 12 (Figure 6C-E). For each cluster, the top 10 differential genes were plotted (Figure 6F). We focused on clusters 0 to 3 containing the majority of the cells (Figure 6E). Clusters 0 and 2 consisted of CD4 T cells while clusters 1 and 3 were CD8 T cells (Figure 6E, S3C, table 1). We therefore labeled these clusters CD4KO, CD8KO, CD4WT and CD8WT. Cluster CD4KO and CD8KO were compared with CD4WT and CD8WT respectively. Over 600 differentially expressed genes were found for each combination (Figure 6G, table 2-3). Interestingly, over 70% of the differentially expressed genes were shared between the CD4 and CD8 populations (Figure S3D). Cluster CD4KO and CD8KO showed high differential expression of *SELL, GZMK, ITGB1, GZMA, CXCR3* while genes associated with terminal exhaustion such as *CCL1, CCL3, CCL4, TNFSRF18, IL2RA, RGS16, SOCS1, TNFRSF9, LAG3, HAVCR2* are strongly downregulated in CD70KO (Figure 6G, table 2-3). We speculated that this unique gene expression pattern could be based on a more exhausted state of the CD70WT nanoCAR T cells. An unbiased dysfunction gene signature of 30 genes that correlated with antigen induced CAR T cell exhaustion^38^ was tested against clusters CD4KO, CD8KO, CD4WT and CD8WT (Figure 6I). As expected, clusters CD4WT and CD8WT highly expressed the dysfunction signature while clusters CD4KO and CD8KO did express this signature to a lesser degree. Furthermore, clusters CD4KO and CD8KO expressed more memory-associated genes such as *TCF7*, *IL7R*, *SELL* and *KLF2* (Figure 6I). Of note, this enrichment was also strongly present in bulk CD70WT and CD70KO nanoCAR T cells (Figure S3E). Exhausted CAR T cells are known to express high amounts of activator protein 1 (AP-1)-related basic leucine zipper (bZIP)–interferon-regulatory factor (IRF) families such as the activation factors JUNB, JUND, FOS, and exhaustion-promoting factors basic leucine zipper ATF-like transcription factor (BATF), BATF3 and IRF4^39^. We observed high expression of these factors in clusters CD4WT and CD8WT while clusters CD4KO and CD8KO, containing mainly CD70KO nanoCAR T cells, showed downregulation of these factors (Figure 6I). Lastly, analysis of the cell cycle revealed an increased frequency of cells within the G2/M and S phases of the cell cycle in the CD70KO condition (Figure 6J). Overall, our single cell analysis showed that CD70KO CD70-specific nanoCAR T cells are less prone to exhaustion. Therefore, we believe that the overall increase of CD70-specific nanoCAR T cell functionality of CD70KO T cells is not so much the result of reduced fratricide but rather due to the prevention of exhaustion.

**Figure 6:**
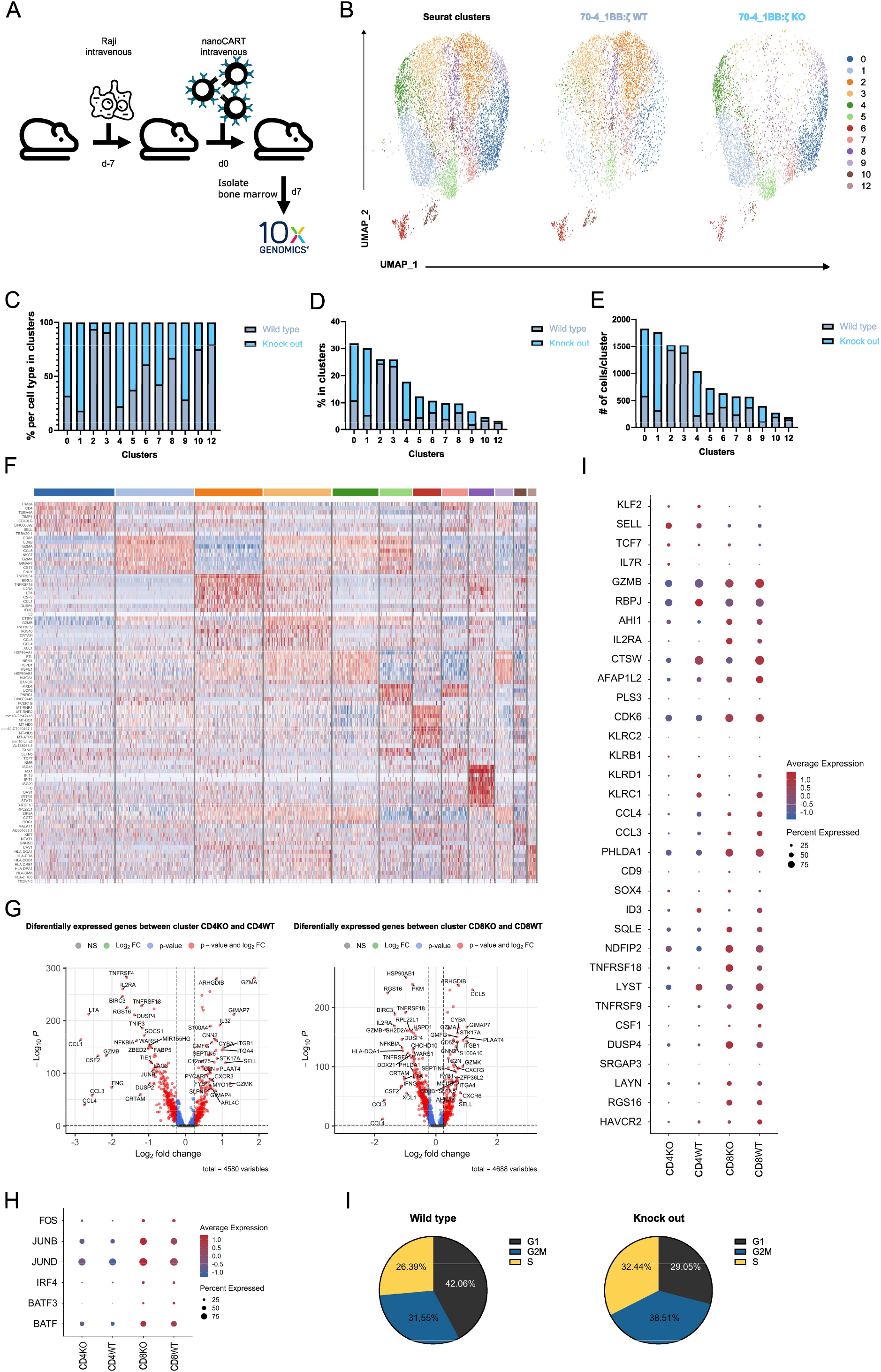
KO CAR T cells are protected from exhaustion. **(A)** The experimental setup of the Raji xenograft model. CD70 nanoCAR T cells (0.75×10e6) were injected intravenously 7 days after intravenous injection of Raji cells (0.5×10e6). At day seven post nanoCAR T cell injection, bone marrow was isolated and cells were sorted for CAR expression. Sorted cells were sequenced after 10x library construction. **(B)** UMAP representation of all the cells with exclusion of cluster 11. UMAP projection of CD70WT (middle) and CD70KO (right) are shown. **(C)** Bar graph representing the proportions of each CAR T cell condition in the different clusters. **(D)** Bar graph representing the contribution of each CAR T cell condition within the different clusters. **(E)** Absolute number of cells per cluster. **(F)** Heatmap of top 10 marker genes for each cluster defined in (B). **(G)** Volcano plot depicting differentially expressed genes between clusters CD4KO and C4WT (0 vs 1) and clusters CD8KO and CD8WT (1 vs 3). Genes upregulated in the KO clusters are on the right side. Red dots indicate significant genes with p < 0.05 and log2FC > 0.2. **(H)** Dot plot illustrating the expression level of naïve/memory genes and the dysfunction gene signature in cluster CD4KO, CD4WT, CD8KO and CD8WT. **(I)** Dot plot illustrating the expression level of *FOS, JUNB, JUND, IRF4, BATF* and *BATF3* in cluster CD4KO, CD4WT, CD8KO and CD8WT. **(J)** Relative frequency of sorted CD70WT and CD70KO nanoCAR T cells in each phase of the cell cycle as determined by single-cell RNA-seq.

## Discussion

In this study, we developed a nanoCAR T cells targeting CD70. First, we showed that JAK/STAT signaling has no beneficial effect on CD70-specific nanoCAR T cells. Surprisingly, the “conventional” 4_1BB-based second generation nanoCAR out performed JAK/STAT signaling nanoCARs. Next, we assessed the functionality in clinically relevant models expressing low to high levels of CD70. The CD70 nanoCAR T cells induced complete elimination of pediatric MRT organoids *in vitro* and improved the survival of DLBCL PDX injected mice *in vivo*. Importantly, we show that CD70 is upregulated on activated nanoCAR T cells and inhibits the function of nanoCAR T cells specific for CD70 by inducing antigen-induced exhaustion. Knocking out endogenous CD70 in CD70-specific nanoCAR T cells resulted in an enhanced tumor eradication, a less-exhausted nanoCAR T cell phenotype with retention of memory phenotype cell populations during *in vivo* stimulation.

CD70 is an attractive target for the treatment of various malignancies. Although its expression is variable within a tumor, we could show here complete elimination both for MRT and DLBCL patient-derived samples. These samples included cells with low to unmeasurable expression. The fact that CAR-T cells were able to eliminate also these cells suggests that even the CD70 negative malignant cells may express the antigen at some point during the rejection phase. In a clinical setting, pharmaceuticals such as azacitdine, which have been proven to increase CD70 antigen density may synergize with CD-70 specific CAR T cell^19,23^. Combining these approaches of boosting antigen density and CAR functionality using the CD70KO, may be an optimized strategy to target AML and solid tumors such as MRT.

Previous reports of CD70KO CD70-specific CAR T cells did not show improved CAR T cell functionality^24^. We believe that the timing of the KO is crucial for the beneficial effect to occur. In contrast to the earlier study, we knocked out CD70 at the earliest stage of T cell activation and before retroviral transduction, resulting in T cells with low to absent CD70 expression before the CAR is expressed. Possibly, continuous CAR stimulation already during the CAR T production process may predispose the T cells for exhaustion. In line with our results, others have targeted membrane proteins expressed by T cells such as CD7 and CD5 either directly by gene-editing or by protein expression blockers (PEBLs) and observed an increase in CAR T cell functionality. This increased functionality was attributed to a decrease in fratricide. In these studies, no assessment of dysfunctionality due to exhaustion was explored^40–42^. Although we did not address the occurrence of fratricide directly, our data suggest that fratricide of nanoCAR^+^ cells is not a major factor contributing to the increased functionality, since the nanoCAR T cells do not express CD70 on the membrane. The absence of CD70 may be due to retention of CD70-CAR complexes in the endoplasmatic reticulum. Alternatively, CD70 may traffic to the membrane and subsequently be removed by interaction with the CAR T cells. This process was named trogocytosis^43^.

We showed that tumor infiltrating CD70WT CD70-specific CAR shown clear signs of exhaustion. CAR T cell functionality is limited by the induction of exhaustion, a progressive state of dysfunctionality that develops when T cells are continuously exposed to antigen and inflammatory stimuli. We hypothesized that the modest functionality of our CD70-specific CAR is the result of increased CD70 mRNA translation upon T cell activation and subsequent binding to and continuous triggering of the CAR, which ultimately results in exhaustion. This exhausition was evidenced by an increased expression of exhaustion-related transcription factors from the bZIP/IRF and AP1 family^39^. CD70WT nanoCAR T cells overexpressed in high amounts *BATF, BATF3, IRF4, NR4A, JUNB* and *JUND*. In addition, we could observe a downregulation of memory associated genes such as *TCF7, IL7R, SELL, CD27, KLF2*. Lastly, a gene set that correlates with exhaustion in CAR T cells was highly enriched in the WT CD70-specific CAR T cells^38^.

Due to the upregulation of CD70 on activated DCs, B and T cells, toxicity can be expected in a clinical setting. It has been shown that CD70-specific CAR T cells could hinder a successful antiviral response^20^. If this is the case, patients treated with CD70-specific CAR T cells could be at risk for infectious diseases caused by CD70-specific immune defects. It is reassuring that multiple groups have shown the safety profile of targeting CD70. For instance, in AML, the first clinical trial with a monoclonal antibody targeted to CD70 in combination with the hypomethylating compound azacitidine showed favorable responses, even complete responses in patients unfit for intensive therapy. Most interestingly, no adverse effects correlated with T cell toxicity were observed^19^. However, CAR T cells are living drugs which make them harder to control compared to a monoclonal antibody and therefore a suicide gene system should be included.

Our results show that CAR T cells targeting CD70 results rapidly in exhausted CAR T cells. Using CRISPR/Cas9 gene-editing technology, we could overcome this exhaustion and significantly increase CAR T cell functionality. These results could influence the optimization of CARs targeting T cell-specific antigens such as CD5, CD4, CD7 and CD30.

## Material and methods

### Culture of cell lines

All the cell lines were cultured as per American Type Collection (ATCC) recommendation. Raji wild type (WT), Raji knock out (KO) and Nalm6 were cultures in Rosewell Park Memorial Institute (RPMI)-1640 medium (Gibco), supplemented with 10% fetal calf serum (FSC) (Biowest), 2 mM L-glutamine (Gibco), 100 IU/mL penicillin (Gibco) and 100 IU/mL streptomycin (Gibco). SKOV3 was cultured in Dulbecco’s Modified Eagle Medium (DMEM) (Gibco, Invitrogen) supplemented with 10% FSC (Biowest), 2mM L-glutamine (Gibco), 100 IU/mL penicillin (Gibco) and 100 IU/mL streptomycin (Gibco).

### Generation of nanoCAR plasmids

The different constructs, as shown in Figure 1A, were generated by Gibson Assembly as described earlier^29^. The CAR backbone constructs were ordered as gBlock (IDT) and cloned into the LZRS-IRES-eGFP retroviral plasmid by Gibson Assembly (NEBuilder HiFi DNA Assembly Master Mix) (NEB). The VHH specific sequence was amplified using PCR (Phusion High Fidelity PCR kit, NEB) and correct overhangs were incorporated. PCR products were visualized on gel and purified using MinElute PCR Purification kit (Qiagen) as per fabricator instructions. The LZRS-CAR-IRES-eGFP plasmids were overnight digested with BamHI (NEB) and purified using Zymoclean Large Fragment DNA Recovery Kit (Zymo) according to manufacture instructions. Subsequently, digested and purified plasmid was dephosphorylated (rAPid Alkaline Phosphatase) (Roche) and used in a Gibson Assembly reaction together with the purified PCR products. The Gibson Assembly reaction mix was transformed in bacteria (NEB Stable Competent Escherichia coli (High Efficiency), NEB) and plated on agar. After overnight incubation, colonies were selected and grown in liquid lysogenic broth (BD Difco) overnight. Plasmids were isolated and sequenced. Colonies containing the correct plasmid were further cultured and midipreps (Qiagen) were performed.

### Generation of retroviral particles

Viral particles were produced as described earlier^29^. In short, retroviral plasmids encoding for the different CAR structures were transfected using calcium phosphate in the amphotropic packing cell line Phoenix A. Every two days, medium was refreshed and every four days cells were selected using puromycin. At day fourteen, retroviral supernatant was collected and frozen at −80°C until use.

### RNP formation

HiFi Cas9 protein was acquired from IDT. CD70 ribonucleoprotein (RNP) was prepared by mixing Cas9 protein and CD70 gRNA (Synthego) at 2.5:1 molar ratio and incubated at 21°C for 10 minutes in a thermocycler and immediately used for nucleofection.

### Generation of CAR expressing T cells

Buffy coats from healthy donors were obtained from the Belgian Red Cross and used following the guidelines of the Medical Ethical Committee of Ghent University Hospital, after informed consent had been obtained, in accordance with the Declaration of Helsinki. Peripheral blood mononuclear cells (PBMC) were isolated by Lymphrep (Axis-Shield) gradient centrifugation. The percentage of CD3^+^ cells was determined by flow cytometry and T cells were stimulated with Immunocult Human CD3/CD28/CD2 T cell activator (StemCell Technologies) per fabricator instructions in IMDM (Gibco) supplemented with 10 % FCS (Biowest), 2 mM l-glutamine (Gibco), 100 IU/mL penicillin (Gibco) and 100 IU/mL streptomycin (Gibco) (complete IMDM, cIMDM), in the presence of 10 ng/mL IL-15 and IL-7 (Milteyni Biotec). After 48 hours, cells were harvested and resuspended in retroviral supernatant and centrifuged for 90 minutes at 1000g at 32°C on retronectin (TaKaRa) coated plates.

For the CD70KO CAR T cells, stimulated PBMC were nucleofected using the Lonza Amaxa 4D-nucleofector X unit (program EH-110). Cells were resuspended at a concentration of 1×10e6 cells/20μl P3 buffer with 20 μM RNP. Following nucleofection, cells were harvested, placed in fresh cIMDM + 10 ng/ml IL-7 + 10 ng/ml IL15 and allowed to rest for 24 hours at 37°C, 7% CO2. After rest, cells were harvested and transduced as described earlier. For the WT CAR T cells, cells were nucleofected but no RNP was added during nucleofection.

### Flowcytometric determination of cytokine production

Detection of cytokine producing cells was performed as previously described. In short, CAR T cells were incubated with target cells for 16 hours in the presence of BD Golgiplug (BD Biosciences). Cells were harvested, labelled with antibodies specific for CD3, CD4 and CD8α and a viability dye was added (Fixable Viability Dye-eFluor 506; Thermo Fisher Scientific). Subsequently cells were fixed and permeabilized (BD Cytofix&Cytoperm, BD Biosciences) following the supplier’s protocol. Cells were analyzed on a LSR II device (BD Biosciences) and data was analyzed using FACS DIVA software (BD Biosciences) and FlowJo v10.8.1 software (BD Biosciences).

### *In vitro* stress test

CAR T cells were added to 96-well plates containing Raji cells at an E:T ratio of 1:1 (1 × 10e4 cells each) in the absence of exogenous cytokines. At the start of the experiment and every two days, wells were harvested and cells were stained for CD3. Before measurement, propidium iodide (Invitrogen) was added. Cells were measured on a LSR II device (BD Biosciences) and data was analyzed using FACS DIVA software (BD Biosciences) and FlowJo v10.8.1 software (BD Biosciences). The remaining wells were stimulated with 10e4 Raji cells.

### *In vivo* experiments

All animal experiments were performed after approval and in accordance with the guidelines of the Ethical Committee for Experimental Animals at the Faculty of Health and Medicine and Health Sciences of Ghent University (ECD 20-18, Ghent, Belgium).

Six to ten week old male or female in-house bred NOD.Cg-Prkdc^scid^IL-2rg^tm1Wjl^/SzJ (NSG) mice were used in all the *in vivo* experiments. Experimental conditions are described within each figure legend. Untransduced T cells, CAR T cells or PBS were injected intravenously via tail vein at the indicated time points. All injected cells were resuspended in PBS and a maximum volume of 200 μl was administered for tail vein injection. For subcutaneous injection, cells were resuspended in 50 μl. Tumor burden was assessed by caliper measurement (SKOV3 model), bioluminescence (Raji model) or tail vein bleeding (DLBCL PDX). Bioluminescence was assessed after intraperitoneal injection of 150 mg/kg bodyweight D-luciferin (Perkin Elmer) using an IVIS Lumina III *in vivo* imaging system. Animals were euthanized as per experimental protocol or when the human endpoints were reached.

### Generation of DLBCL PDX

The DLBCL PDX model (LYM024) was derived at first relapse after R-CHOP in a patient with a DLBCL, non-GC subtype by IHC, BLC2 and cMYC double-expressor, without BCL2 or cMYC rearrangements with negative CISH EBV (EBER). It was obtained from the registered biobank “PDX Hematologie UZGent” (Belgian Registration number BB190094). Tumor cells were isolated from early passage (P2) tumor bearing NSG mice and intravenously injected in FBS in new NSG mice.

### Patient-derived MRT organoid culture

MRT organoids were established, characterized, and cultured as previously described^35,44^. In brief, MRT organoids were cultured in droplets of growth factor-reduced basement membrane extract (BME) Type 2 (R&D Systems) in kidney organoid medium (KOM) (Advanced DMEM/F12 (Gibco) containing 1X GlutaMAX (Thermo Fisher Scientific), 10 mM HEPES (Thermo Fisher Scientific) and 1X Penicillin-Streptomycin (P/S) (Merck Millipore) (AdDF+++), supplemented with 10% R-spondin–conditioned medium, 1.5% B27 supplement (Gibco), 50 ng/mL epidermal growth factor (EGF) (PeproTech), 50 ng/mL fibroblast growth factor (FGF-)2 (PeproTech), 1.25 mM N-acetylcysteine (Sigma), 10 μM Rho-associated coiled–coil containing protein kinase (ROCK) inhibitor Y-27632 (Abmole) and 5 μM A83-01 (Tocris Bioscience)). Culture medium was refreshed every 3 to 4 days and organoids were dissociated mechanically once a week.

### Bulk RNA sequencing analysis

Bulk RNA sequencing data from pediatric renal tumor tissue and organoids as well as normal kidney organoids were used to determine RNA expression of CD70^45^.

### Flow cytometry analysis

Staining of surface markers was performed in DPBS (Gibco) with 1 % FCS (Biowest) using the antibody to cell ratio recommended by the supplier.

For all samples, after antibody incubation cells were washed once with FACS buffer and propidium iodide (Invitrogen) was added. Flow cytometric analysis was performed on a LSR II device (BD Biosciences) and data was analysed using FACS DIVA software (BD Biosciences) and FlowJo v10.8.1 software (BD Biosciences).

#### Staining on MRT organoid cultures

To determine CD70 expression on MRT organoids, organoids were mechanically dissociated into a single cell suspension, washed twice, and resuspended in FACS buffer (PBS, 2% FCS, 2 mM EDTA). Cells were stained with anti-CD70 (Cusatuzumab, MedChemExpress) labeled with Alexa Fluor™ 647 NHS Succinimidyl Ester (Invitrogen) in a 1:1000 dilution for 30 min on ice.

CD137 expression on CAR T cells was determined after 20 hours of co-culture with MRT organoids. Cells were mechanically dissociated, washed twice, and resuspended in FACS buffer. Cells were stained with CD137-APC (BD Biosciences) in a 1:25 dilution for 25 min on ice.

For all samples (during MRT organoid cultures), after antibody incubation cells were washed three times with FACS buffer and DAPI was added in a 1:200 dilution as a viability dye. Surface antigen expression was measured with the CytoFLEX LX (Beckman Coulter) and data was analyzed with the CytExpert Software Version 2.4 (Beckman Coulter) and FlowJo v10.8.1 software (BD Bioscienes).

### Co-culture conditions

MRT organoids were mechanically dissociated four days before the co-culture assay and seeded out in BME droplets in KOM. One day prior to the co-culture 70- and 20-4_1BBζ cells were defrosted and cultured in a density of 1 × 10^6 cells/mL in RPMI containing1X GlutaMAX (Gibco), 1X P/S and 10% FCS supplemented with 10 ng/mL IL7 and IL15 overnight. On the day of co-culture, MRT organoids were carefully collected, washed with cold medium to remove BME, and a fraction was dissociated into a single cell suspension. Single tumor cells were counted to determine the number of cells present in the MRT organoid suspension. MRT organoids were then seeded out in suspension in co-culture medium containing 10% FBS and a 1:1 ratio of KOM and RPMI with 1X GlutaMAX and 1X P/S. Next, CAR T cells were washed, counted and according to the chosen E:T ratio an appropriate amount was added to the MRT organoids in co-culture medium. The cells were incubated for the indicated time points and medium was refreshed on day 2 for the 3-day co-culture and on day 3 and 6 for the 7-day co-culture.

### MRT organoid luciferase killing assay

To determine tumor cell specific killing, a lentiviral transduction with pLKO.1-UbC-luciferase-blast was performed on all MRT organoid lines, as described previously^46–48^. Two days after transduction, luciferase-transduced cells were selected by addition of 10 μg/mL blasticidin. For the co-culture assay, luciferase expressing MRT organoids were seeded in a flat-bottom 96-well plate with an equivalent of 7500 single tumor cells per well and CAR T cells were added in the indicated ratio in total amount of 100 μL of co-culture medium. Luciferase activity was determined after washing the cells with PBS, using the luciferase assay system with the Passive Lysis 5X buffer (Promega) according to manufacturer’s instructions.

### Live cell imaging

Live cell imaging was performed as described previously with minor modifications^49^. In brief, to visualize 70- and 20-4_1BBζ CAR T cells in the co-culture assay, cells were washed with PBS and stained with eBioscience Cell Proliferation Dye eFluor 450 (Thermo Fisher Scientific) in PBS in a 1:1000 dilution for 10 minutes at 37°C. RPMI with 10% FBS was added, and cells were further incubated on ice for 5 minutes. Then cells were washed, counted and seeded out in black, glass-bottom 96-well plates (Greiner) together with MRT organoids in the indicated ratio. The co-culture medium was supplemented with 2 drops/mL of NucRed Dead 647 ReadyProbes Reagent (Thermo Fisher Scientific) to fluorescently label live (excited by 561 nm laser) and dead cells (excited by 633 nm laser). Cells were incubated in the incubation chamber of the Leica TCS SP8 confocal microscope (5% CO2, 37°C) and imaged every 30 minutes with a HC PL APO CS2 10x/0.4 dry objective for 72 hours. The following settings were used: resonant scanner at 8000 Hz, bidirectional scanning, line averaging of 8, optimal Z-stack step size, Z-stacks of 60-110 μm in total and a resolution of 512 × 512 resulting in a Voxtel size of 1.82 μm × 1.82 μm × 2.409 μm.

### scRNA-seq

#### Single cell library preparation and sequencing

Mice were injected with 5×10e5 Raji cells. After seven days, CAR T cells were intravenously injected. Mice were sacrificed seven days post CAR T cell injection. Bone marrow was isolated and processed to obtain a single cell suspension. Single cell suspension was labelled with antibodies for human CD45, CD19 and hashtag antibodies. The suspension was sorted on a BD ARIA II device based on positivity for human CD45, human CD19 and eGFP. Sorted cells were pooled and resuspended in PBS + 0.04% BSA to obtain on average 1200 cells/μl and loaded on a Chromium GemCode Single Cell Instrument (10x Genomics) to generate single-cell gel beads-in-emulsion (GEM) at the VIB Single Cell Core. The scRNA-Seq libraries were prepared using the GemCode Single Cell 5’ Gel Bead and Library kit, version NextGEM 2 (10x Genomics) according to the manufacturer’s instructions. Sequencing libraries were loaded on an Illumina NovaSeq flow cell at VIB Nucleomics core with sequencing settings according to the recommendations of 10x Genomics.

#### Processing of scRNA-seq and hashtag data

The Cell Ranger pipeline (10x Genomics, version 6.0.0) was used to perform sample demultiplexing and generation of FASTQ files for read 1, read 2 and the i7 sample index for the gene expression and hashtag libraries. Read 2 of the gene expression library was subsequently mapped to a combined reference genome of hg38 and mm10. The resulting count matrices were loaded into R and processed using the Seurat package (version 4.0.5)^50^. The hashtag data was normalized via the “NormalizeData” function using the CLR (Centered Log Ratio) method. Subsequently the “HTODemux” function was used to demultiplex the different cell types i.e., WT and KO. Therefore, the positive.quantile parameter was set to 0.99. After demultiplexing, only the singlet cells were retained for further analysis.

The singlet cells were filtered further based on their gene expression data using the following criteria: i) nFeature_RNA > 600; ii) nCount_RNA > 1143; iii) nCount_RNA < 30000 and iv) percent.mt < 10. Prior to further processing of the gene expression data, three gene sets were removed, namely Cell Cycling and Histone genes^51^ and CD70. For removal of the Cell Cycle genes, we relied on Table S8 of Park *et al*.^52^. Afterwards the gene expression data was normalized with the “NormalizeData” function using the LogNormalize method and a scale.factor of 10000. Then the highly variable features were identified with the “FindVariableFeatures” function with selection.method set to vst and nfeatures to 4000. After which the data was scaled with the “ScaleData” function.

For visualization of the data the “RunPCA” and the “RunUMAP” functions were used. After which clustering was performed using the “FindNeighbors” and “FindClusters” functions. Differential expression analysis was performed via the “FindMarkers” function for all clusters, for comparison of the two cell types i.e., KO and WT and for comparison of specific clusters.

### Statistical analysis

In general, data is shown as mean ± standard error of the mean, unless otherwise noted in the figure legend. Significance was considered at p < 0.05. All statistical tests are described in the figure legends and p values are denoted either exact or with asterisk as follows: **** = p < 0.0001; *** = p < 0.001: ** = p < 0.01; * = p < 0.05. Number of replicates are described within the relevant figure legend. Data was analysed and visualized using GraphPad Prism v9.5.0 (Dotmatics).

## Acknowledgements

We thank the facilities of the Princess Máxima Center for imaging, organoid culture as well as the flow cytometry for support. We thank C. Janda (Princess Máxima Center) for providing the anti-CD70 antibody labeled with Alexa Fluor™ 647. We are grateful for the support of the Oncode Institute and the DFG. Further, we thank Prof. Dr. Tom Boterberg (Ghent University Hospital) for the irradiation of cells and Prof. Dr. Kevin Braeckman (Ghent University) for provision of the nucleofector.

## Author contributions

Conceptualization: SDM, BVDK; Methodology: SDM, BVDK, JB, JD; Software: LDC, NV; Validation: BVDK, JD; Formal analysis: SDM, LDC; Investigation: SDM, JB, AVP, GG; Resources: SDM, JB, WD, ED, JT, JD, BVDK; Data curation: SDM, LDC, JB, BVDK; Writing – original draft: SDM, BVDK; Writing – review & editing: SDM, JB, LDC, AVP, WD, EP, LD, JI, GG, LB, MP, NV, JVD, FO, TK, ED, JT, JD, BVDK; Visualization: SDM, JB, LDC; Supervision: TK, JB, BVDK; Project administration: SDM, BVDK; Funding acquisition: JD, BVDK.

