## Supplemental figures for "Knocking out CD70 rescues CD70-specific nanoCAR T cells from antigen induced exhaustion"

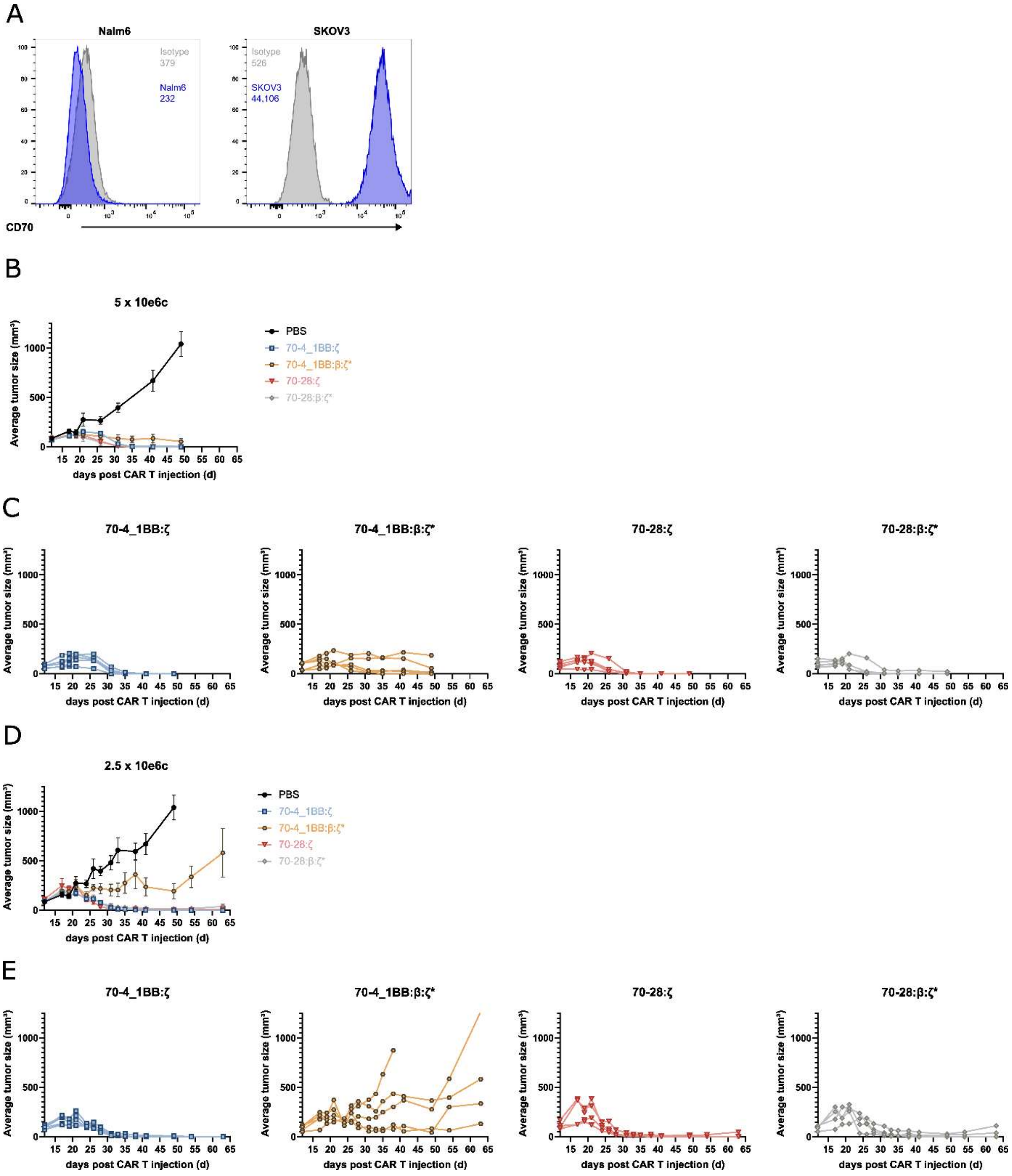


**Figure S1: CD70-specific CAR T cells eliminate SKOV3 subcutaneous tumors at higher CAR T cell doses. (A)** Expression of CD70 on NALM6 and SKOV3 cell line. MFI is indicated, grey = isotype, blue = CD70-specific monoclonal antibody. **(B)** SKOV3 tumor growth after injection of CAR T cells (5 x 10e6). Tumor growth is shown as average tumor size, error bars indicate SEM. **(C)** Tumor growth for each individual mouse. **(D)** SKOV3 tumor growth after injection of CAR T cells (2.5 x 10e6). Tumor growth is shown as average tumor size, error bars indicate SEM. **(E)** Tumor growth for each individual mouse.

A

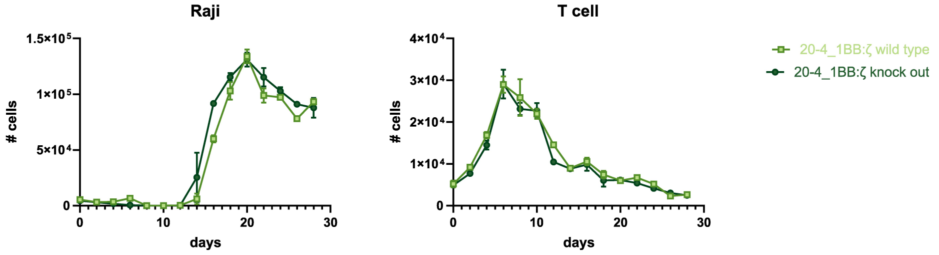


**Figure S2: CD70KO has no effect on CD20-specific nanoCAR T cells. (A)** CD20 specific nanoCAR T cells either WT or KO were incubated with WT Raji cells. Every two days the amount of cells were determined and the remaining wells were stimulated with fresh Raji cells. Median values are shown, error bars indicate SEM.

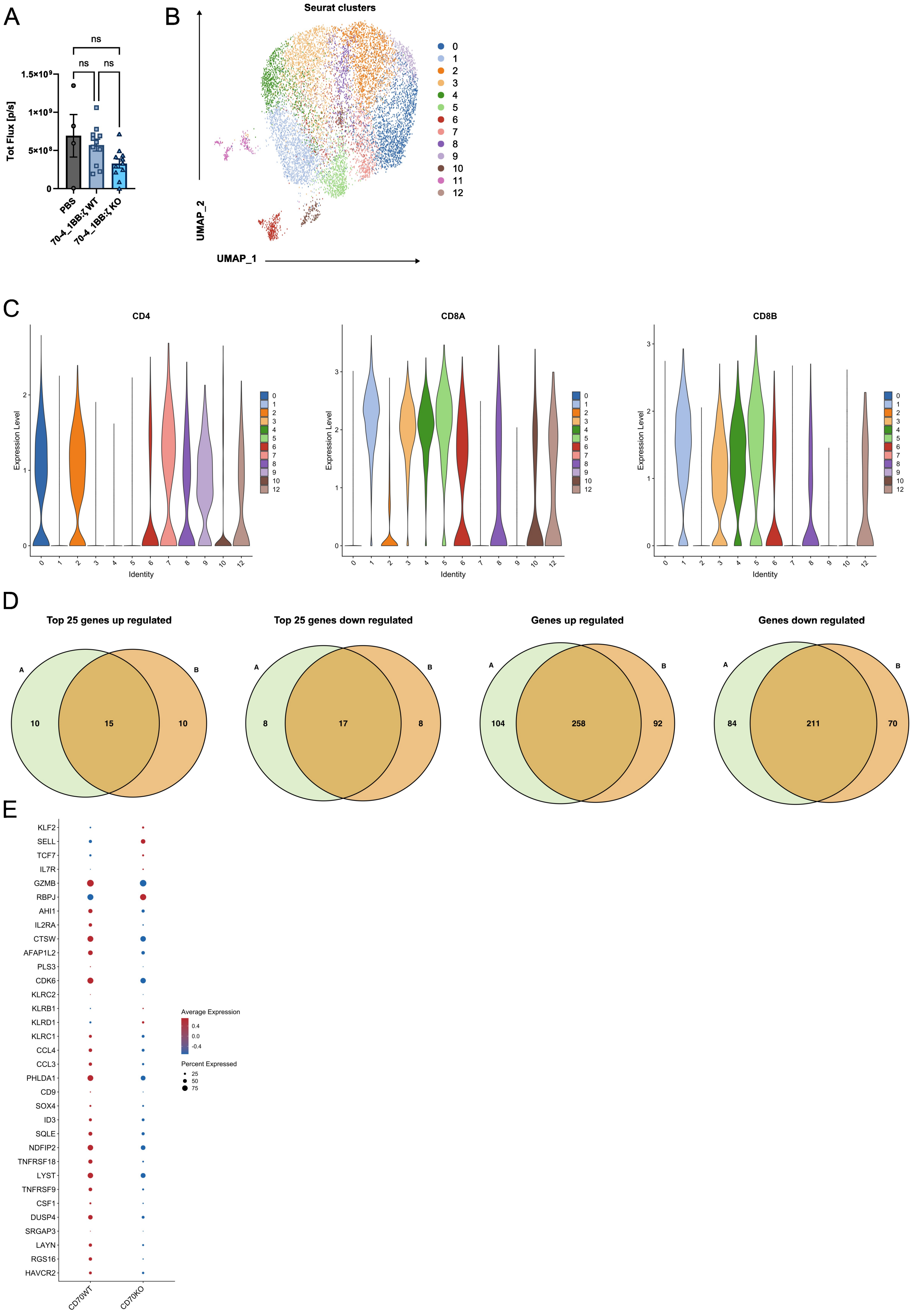


**Figure S3: CD70 KO protects CD70-specific CAR T cells from exhaustion. (A)** Presence of Raji cells in individual mice measured as total flux (photons/s). **(B)** UMAP representation of all the cells. **(C)** Expression of *CD4*, *CD8A* and *CD8B* for each cluster, excluding cluster 11. **(D)** Venn diagrams showing overlap of genes between the differential expression analysis of CD4KO vs CD4WT and CD8KO vs CD8WT. **(E)** Dot plot illustrating the expression level of naïve/memory genes and the dysfunction gene signature in WT and CD70 KO CAR T cells.
